# A kiwellin protein-like fold containing rust effector protein localizes to chloroplast and suppress cell death in plants

**DOI:** 10.1101/2021.08.20.456821

**Authors:** Rajdeep Jaswal, Sivasubramanian Rajarammohan, Himanshu Dubey, Kanti Kiran, Hukam Rawal, Humira Sonah, Rupesh Deshmukh, Tilak Raj Sharma

## Abstract

The effector proteins expressed by plant pathogens are one of the essential components of the host-pathogen interaction. Despite being important, most of the effector proteins remain unexplored due to the lack of conserved features and huge diversity in their primary sequence. In the present study, extensive secretome analysis was performed in sixteen major plant fungal pathogens to find the conserved features in the candidate secretory effector proteins (CSEPs) using homology and *ab initio* modeling approaches. Interestingly, a variable number of plant kiwellin proteins fold like secretory proteins were found in all the major rust fungal pathogens. Many of them are predicted as potential effector proteins. For instance, 26 out of 35 Kiwellin like proteins identified in *Puccinia striiformis race 104E 137A* were predicted as potential effector proteins. In addition, a kiwellin predicted effector gene, *Pst_13960*, from the Indian *Puccinia striiformis race Yr9* was characterized using overexpression, localization, and deletion studies in *Nicotiana benthamiana*. The *Pst_13960* suppressed the BAX-induced cell death and localized in the chloroplast. Furthermore, the expression of the kiwellin matching region (*Pst_13960_kiwi*) alone suppressed the BAX-induced cell death in *N. benthamiana* despite the change of location to the cytoplasm and nucleus, suggesting the novel function of the kiwellin fold in rust fungi. Further analysis of these proteins predicted these candidates to contain N-terminal Intrinsically disordered regions (IDRs) putatively associated with chloroplast translocation as deletion of region abolished the chloroplast localization of *Pstr_13960*. Overall, the current study reports the presence of kiwellin like proteins in rust fungi that act as a novel effector in plants.

**Author Summary:** Rust fungi are one of the most devastating plants infecting pathogens. These pathogens secrete several distinct proteins like effector proteins that help the pathogens in the establishment of infection by suppressing cell death induced by the plants. Despite being important, these effector proteins remain unexplored due to the lack of conserved features. Currently, different methods are being used to characterize them however, could not describe their specific function fully due to a lack of knowledge of the functional domain. Recent advancement in effector protein tertiary structure characterization using NMR (Nuclear magnetic resonance) and X-ray crystallography has been very helpful in identifying the conserved structural features defining functionality. However, these techniques are quite complicated and may take a lot of time and labor. On the other hand, the computational approaches for structural prediction of the effectors may help to identify known folds or domains with few efforts but at a significant level. Therefore, such computational approaches can be efficiently implemented in the preliminary screening of the candidates. In the present study using the computational structure prediction method, we were able to find several conserved novel kiwellin folds containing effectors, in different rust fungi. We characterized one of the candidates and it showed interference with artificially induced cell death in plants. This study highlights the novel function of the kiwellin like effector proteins of the rust fungi that are already identified to play a role in host defense against plant pathogens.

## Introduction

All plant fungal pathogens use secretome, and their independent entities, small secretory proteins, explicitly known as effectors as an essential weapon to defeat the host resistance mechanism irrespective of nutritional style. It is now evident through various studies that understanding the secretome can provide the solution to reduces losses due to crop diseases. Moreover, with the commencements of genome sequencing studies, unraveling the pathogen secretome has become very much feasible. A large number of economically important necrotrophs, hemibiotrophs, and biotrophs have been studied to elucidate the secretomic differences and further potential avirulence candidates to develop resistant varieties[1,2]. However, irrespective of large genomic resources and various computational tools, the complete understanding of the pathogen secretome is accompanied by several hurdles. The major problem in understanding the secretome is the absence of conserved features among the effector proteins, especially for obligate biotrophic pathogens [1,3]. The secretome of the necrotrophs and hemibiotrophs is comparatively less equipped with the novel effector proteins as their lifestyle involves necrosis and cell death than manipulations of host metabolism.

Among the obligate biotrophs, the rusts are among the most destructive biotrophic pathogens of the class basidiomycetes [4,5]. These pathogens are macrocyclic and heteroecious in their lifecycle; they are also highly host-specific or, in several cases, species-specific, further causing them to be unique in secretome consortium [6,7]. The rusts are known to encode the largest genome size, a high number of secretory proteins, and species-specific proteins[8,9]. These species-specific proteins are mostly novel and do not show any similarity to known domain proteins at the sequence level[10,11]. To maintain the biotrophic lifestyle and manipulate host defense simultaneously, the rust deploys a plethora of effector proteins. However, in resistant varieties of hosts, the host immunity genes impose selection pressure, causing these effectors to change their gene sequences to further avoid the recognition of their protein product continuously by the host resistance proteins [12,13]. The selection pressure changes the effector sequence; however, to continue their native functions, these proteins may keep their conserved fold rather than maintaining the full domain[14]. Several studies have recently revealed the assumptions of maintaining the conserved molecular function of the effector due to their conserved fold at 3 Dimensional level despite no sequence similarity at the sequence level [14]. The structural genomics of two poplar rust effectors, *MLP124266* and *MLP124017*, using nuclear magnetic resonance (NMR), showed similarity to knottin and nuclear transport factor 2-like proteins[15].

Similarly, the effectors family that includes *AVR1-CO39* (*Magnnaporthe oryzae*), *AvrPiz-t* (*M. oryzae*), and *ToxB* effectors (*Pyrenophora tritici-repentis*) [14], and several other effectors present in many ascomycetes pathogens that do not share similarity among themselves at the sequence level; however, possess a conserved six β-sandwich structure, stabilized by a disulfide bridge between 2 conserved cysteines residues [16,17]. Similarly, several other effectors from *Blumeria graminis* are known to exhibit similarity to ribonucleases (RNase-like effectors) and are involved in the host manipulation[18]. These studies have been clearly shown that the arms race has led them to diverge the primary sequence of effectors; simultaneously, functional necessity has sustained these proteins to keep conserved folds[14,18].

Kiwellins proteins are members of a cysteine-rich major allergen protein family, present in kiwi fruits and reported to cause allergic reactions in humans after consumption[19]. In addition to their role as an allergen in kiwi plants, no designated role has been assigned to those proteins until Han et al., (2019), proposed their role in the host defense against plant pathogens[20–22]. Homologous of the kiwellins are also reported in all major plant species, including conifers and angiosperms, and several cases are reported to be present in high members suggesting the expansion of this family[20]. The study by Han et al., (2019) defined these proteins to block the functioning of one fungal effector, *Cmu1*, known to down-regulate the salicylic acid synthesis by channeling its substrate to other pathways [20–23]. Han et al. (2019) showed that *ZmKWL1*, one of the Kiwellin from the 20 encoded maize Kiwellin, binds to *the Cmu1* active site so that *Cmu1* could not bind to its substrate or dimerize with host Chorismate mutase. The study also defined that plant Kiwellin possesses unique folds that specifically bind to fungal *Cmu1* rather than their native Chorismate mutase (CMs). Han et al. (2019) has also identified kiwellin protein in fungal pathogens; however, to our knowledge, no Candidate secretory effector proteins (CSEPs) have been reported to possess kiwellin like folds in plant-pathogen effector proteins.

Here we report novel kiwellin fold containing effectors using structural modeling in different rust fungi. Additionally, using refined analysis we showed that DPBB domain-containing proteins in the rust secretome and other plant pathogens may possess kiwellin proteins like fold. However, rust fungi may have improvised themselves to use these fold-containing proteins as effectors as none of the analyzed other plant-pathogen showed secretory or effector features in the identified kiwellin fold proteins. The analysis of the rusts proteins also revealed them to possess N-terminal IDRs. Further deletion fragments of *Pstr_13960* revealed the N-terminal and IDR region putatively responsible for the chloroplastic targeting of *Pstr_13960*. Moreover, we also showed that the kiwellin matching region of *Pstr_13960* alone is sufficient to suppress BAX-induced cell death in *N. benthamiana* despite its changed location to cytoplasm and nucleus. Further, the docking analysis of host chorismate proteins (CMs) with rusts kiwellin effectors showed the positive interaction of these proteins suggesting that the rust effector may capture the host CMs as an alternate strategy to block salicylic acid synthesis. The present work is the first study to report the presence of kiwellin like-fold effectors (KLEs) in rust fungi and their expansion in their secretome. The study further highlights the use of the KLEs to target chloroplast and exhibiting cell death suppression possibly binding by plant chorismate mutase.

## Materials and methods

### Secretome analysis and identification of novel CSEPs in the proteome of plant pathogens

The predicted proteome data of plant fungal pathogen was retrieved from Ensembl fungi, except for *Puccinia triticina*, *P. striiformis*, that was utilized from Kiran et al. [10,11]. The genome of *P. arachidis* and *P. horiana* was downloaded from the NCBI database, and genes were predicted using MolQuest 2.0 (FGENESH). The secretome analysis was carried using the secretool online server with default parameters [24]. The conserved domain analysis was done using the Pfam and NCBI-CD Batch search facility using default parameters[25,26] The candidate proteins with no hits were treated as novel secretory proteins. These proteins were further analyzed using EffectorP 2.0 software for potential effector candidates [27]. The proteins with scores above 0.5 were treated as potential candidate effectors.

### Structural modeling of CSEPs and docking

The homology modeling of the CSEPs was done using PHYRE2 Protein Fold Recognition Server, Robetta, and *ab initio* using I-TASSER[28–30]. The 3D structure with template similarity of a minimum of 20% and the confidence score of > 60% proteins were counted as significant results. The protein hits with less than the threshold were removed from the analysis. The significantly modeled proteins were also manually crosschecked for the presence of any conserved domain using NCBI-CD search if any. The Ramachandran plot and model stability were checked using the SAVES server (https://saves.mbi.ucla.edu/). Motif analysis was done using the MEME motif at default settings(https://meme-suite.org/meme/doc/cite.html). Docking analysis was performed using default parameters using FRODOCK[31].

### Identification, sequence alignment, and motif analysis of kiwellin protein fold containing candidate rust effectors

The sequence alignment of full-length proteins and the conserved region was done using MAFFT software[32]. For comparative analysis, the plant kiwellin proteins were downloaded from NCBI. To identify Kiwellin fold-like proteins in other plant pathogens and other rust races, we performed genome-wide identification using randomly selected rust kiwellin and plant kiwellin proteins against NCBI nr database and MycoCosm fungi genome database. The significantly matched proteins (e-5) were selected for structural analysis.

### Gene structure and Phylogenetic analysis

The intron-exon boundaries were predicted using the GSDS server (http://gsds.gao-lab.org/). The Mafft software was used for aligning the rust fungi-specific Kiwellin like effector proteins, and an aligned file was used to generate a maximum likelihood method-based phylogenetic tree using the IQ-TREE server (http://iqtree.cibiv.univie.ac.at/).

### Disordered region prediction and chloroplast prediction

The disordered region was predicted using, SLIDER, PONDER, IUpred2A tools using default parameters[33–35]. For chloroplast transient peptide was predicted using ChloroP, Localizer, and WolfPsort[36–38].

### Cloning of candidate rust KLE genes

The CDS of *Pstr_13960* (with and without signal peptide coding sequence) was PCR amplified and cloned in pENTR/Dtopo vector followed by confirmation with restriction and sequencing. The confirmed *pENTR_13960* plasmid clone was further mobilized in pGWB408 (overexpression) and pGWB441 (subcellular localization) using LR clonase enzyme mix (Invitrogen, USA). The kiwellin domain matching region was cloned separately for overexpression and localization studies similarly as mentioned for full length *Pstr_13960*. Full length CDS of the BAX gene was PCR amplified from mouse cDNA and further cloned in the pGWB408 vector.

### Strains plant material and growth conditions

*Escherichia coli* Topo10 cells (Invitrogen, USA) were used for cloning all the genes. About 4 weeks old *N. benthamiana* plants grown at 22□°C with a 14□hours light/10□hours dark cycle were used for agroinfiltration assays. The *Agrobacterium tumefaciens* strain GV3101 chemically competent cells were transformed with *Pstr_13960*_pGWB441 (subcellular localization) and *Pstr_13960*_pGWB408 (overexpression) constructs. The resulting transformants were further confirmed by PCR.

### Agroinfiltration, subcellular localization overexpression, and phenotype analysis

The confirmed agrobacterium containing localization and overexpression constructs clones were overnight grown (28□°C) and adjusted to an optical density (OD) of 0.4, centrifuged for 10□minutes at 4,000× g. The pellet was re-suspended in MES-K buffer (100μM acetosyringone, 10□mM MgCl2, and 10□mM MES, pH 5.6) and further incubated five hrs at room temperature before infiltration on *N. benthamiana* leaves. For cell death, masking analysis, the candidates genes were infiltrated to *N. benthamiana* leaves 24 hrs later, BAX infiltration was done at the same spots on the leaves.

## Results

### Secretome analysis of plant pathogens and identification of unannotated proteins

The secretome analysis of plant pathogens belonging to different lifestyles revealed a variable number of secretory proteins encoded in their genome. The highest secretory proteins were found in the *Magnaporthe oryzae*, followed by *Fusarium oxysporum* and *Puccinia striiformis Yr9. In* contrast to this, the lowest numbers of secretory proteins were found in *Puccinia psidii* (140) and *Blumeria graminis* genome (193) (Table 1). The identification of unannotated proteins from the secretome also showed *P. sojae* to encode the largest (410) unknown novel domain-containing proteins followed by *Puccinia triticina 77-5* encoding 409 novel proteins. The lowest unannotated proteins were found in necrotroph *Rhizoctonia solani* (119).

**Table 1.** Identification and analysis of the secretome of various plant pathogens.

| Species | Class | Mode of nutrition | Secretory proteins | Unannotated proteins |
| --- | --- | --- | --- | --- |
| *Ustilago maydis* | Basidiomycetes | Biotroph | 273 | 134 |
| *Blumeria graminis* | Ascomycetes | Obligate biotroph | 193 | 120 |
| *Rhizoctonia solani* | Basidiomycetes | Necrotroph | 361 | 119 |
| *Botrytis cinerea* | Ascomycetes | Necrotroph | 432 | 147 |
| *Fusarium oxysporum* | Ascomycetes | Hemibiotroph | 591 | 205 |
| *Magnaporthe oryzae* | Ascomycetes | Hemibiotroph | 639 | 305 |
| *Puccinia triticina* 77-5 | Basidiomycetes | Obligate biotroph | 534 | 409 |
| *Puccinia graminis* 40-3 | Basidiomycetes | Obligate Biotroph | 405 | 300 |
| *Puccinia striiformis* Yr9 | Basidiomycetes | Obligate biotroph | 557 | 407 |
| *Phytophthora infestans* | Oomycetes | Hemibiotroph | 671 | 265 |
| *Phytophthora sojae* | Oomycetes | Hemibiotroph | 1030 | 410 |
| *Pythium ultimum* | Oomycetes | Necrotroph | 561 | 175 |

### Rust fungal pathogen encodes a large number of KLEs proteins in their secretome

The pipeline used for identifying novel secretory effector proteins is shown in Figure.1. The structurally modeling of the novel secretory proteins for most of the studied species revealed non-significant similarities to several proteins. Surprisingly, in the case of rust fungi, we found several candidates matching with kiwellin proteins that are present in eukaryotes especially plants (Supplementary file S1) (modeling parameter, >20% similarity, > 60 % modeling confidence). The results generated by PHYRE2 were also confirmed by I-TASSER and Robetta for randomly selected proteins and these softwares also predicted these proteins to match with plant kiwellin proteins. Thereafter, to further explore kiwellin proteins’ status in all major rust pathogens, these proteins were mined across the genome (Table1). The analysis returned a variable number of proteins similar to kiwellin proteins (minimum 1 and maximum 35 per rust species) (Table 1) Further, the NCBI CD-search revealed these proteins to possess DPBB at the sequence level (Supplementary file S2). However, the structural analysis revealed that most of these proteins matching with plant kiwellin protein fold(Supplementary file S3). As the DPBB fold is present in several other proteins such as endolytic peptidoglycan transglycosylase RlpA, EG45-like domain proteins, plant kiwellins and expansins, bacterial papain inhibitor, and expansins. We designated only those proteins with kiwellin containing folds when the top 3 hits of modeled protein matched with Plant Kiwellins and Ripening-related proteins of plants. The analysis of significantly matching candidate kiwellin like proteins for secretory and effector features revealed most of these were predicted to be secretory (1 to a maximum of 28 proteins per rust species genome) and candidate effectors using EffectorP 2.0 (1 and maximum 26 proteins in rust species genome) (Table 1) The structural analysis also revealed these proteins to have variable similarity and modeling scores when compared to plant kiwellins. The highest sequence similarity (45%) was demonstrated by PLW22586.1 and PLW05486.1 (*P. coronata*) and modeling confidence (97.8%) by *Pstr_13960*, *Pucstr1_11*799, and *Pucstr1_5472* all belonging *to P. striiformis* only. (Supplementary files S1 and S3).

**Fig 1.**
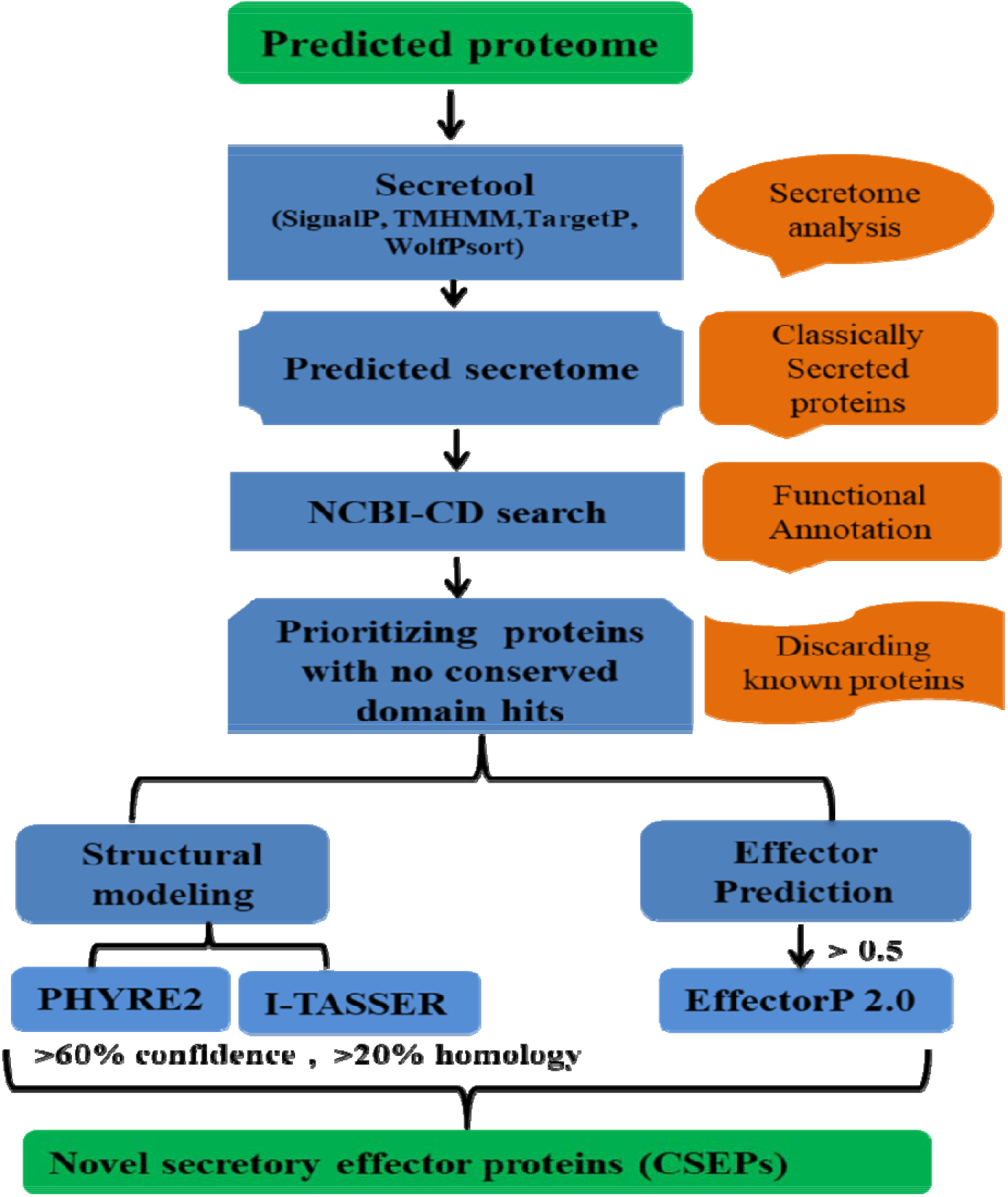
Secretome and annotation pipeline for identification of novel candidate secretory effector proteins (CSEPs).

### Predicted Kiwellin like-fold of rust effectors significantly align with the crystal structure of plant kiwellin proteins

Out of 109 modeled rust KLE candidates the structural analysis of the randomly selected candidates from different rust species showed stable 3D structure as predicted by the Ramachandran plot and SAVES server analysis. Further, the structural analysis showed all these proteins to possess a central β-barrel domain composed of anti-parallel β-sheet similar to the crystal structure of maize (Zea mays, PDB Id 6TI2) and kiwi kiwellin proteins (*Actinidia chinensis*, PDB Id-4PMK) (Figure 2 and 3). Further, the superimposition of the rust KLEs with 4PMK and 6TI2 showed significant superimposition with RMSD in the range of 1-2 and TM score >0.5 thus showing conserved DPBB and Kiwellin protein-like structure (Figure 4).

**Fig. 2.**
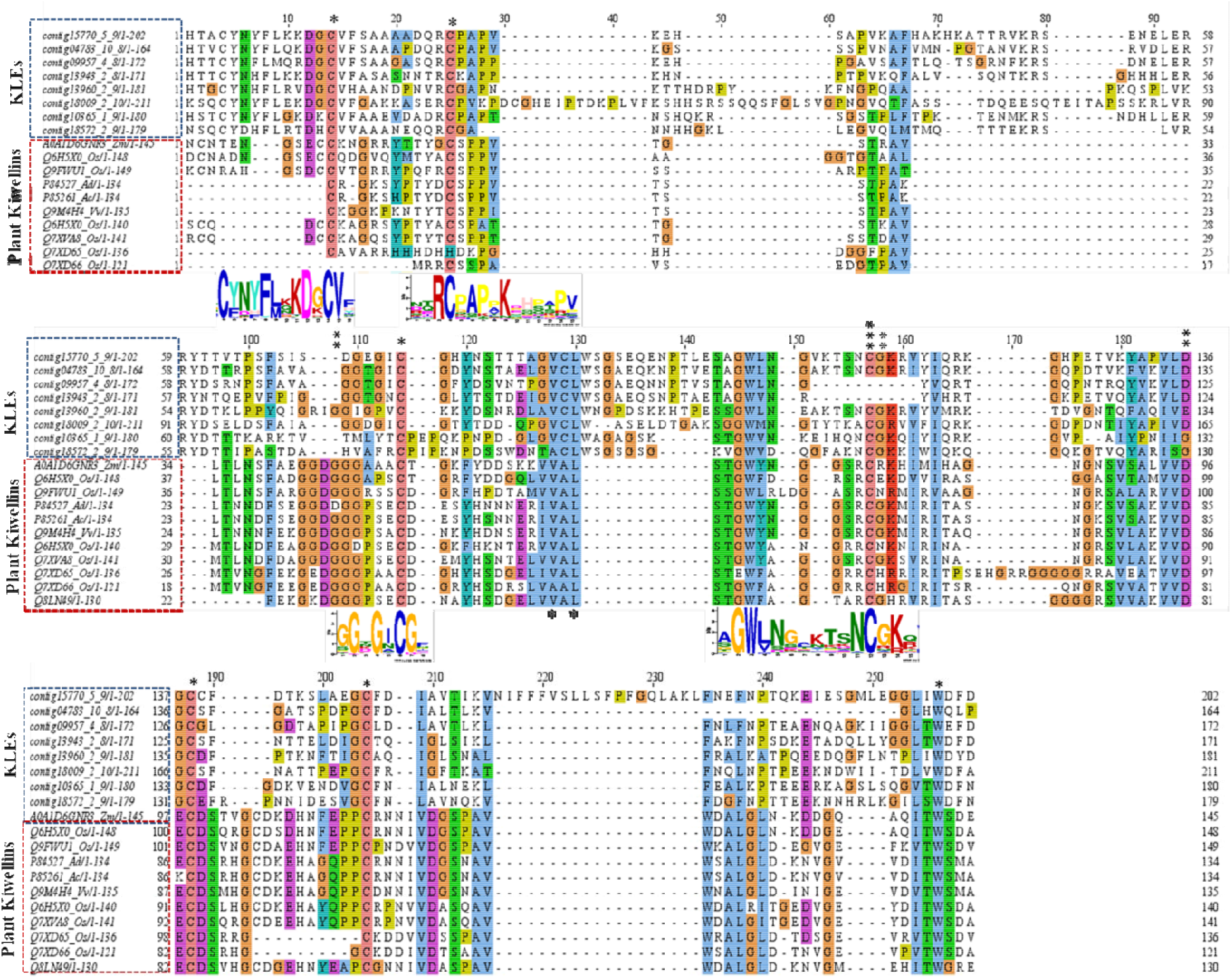
Sequence analysis showing the conserved region of plant kiwellin proteins with selected rust KLE proteins. The residues with a single asterisk sign are highly conserved in rust KLEs and plant kiwellin proteins. The double asterisk represents residues involves in KWl or KWLb interaction to the Cmu1 effector whereas the triple asterisk highlights residues used by KWl1 and KWLb for *Cmu1* binding.

**Fig 3.**
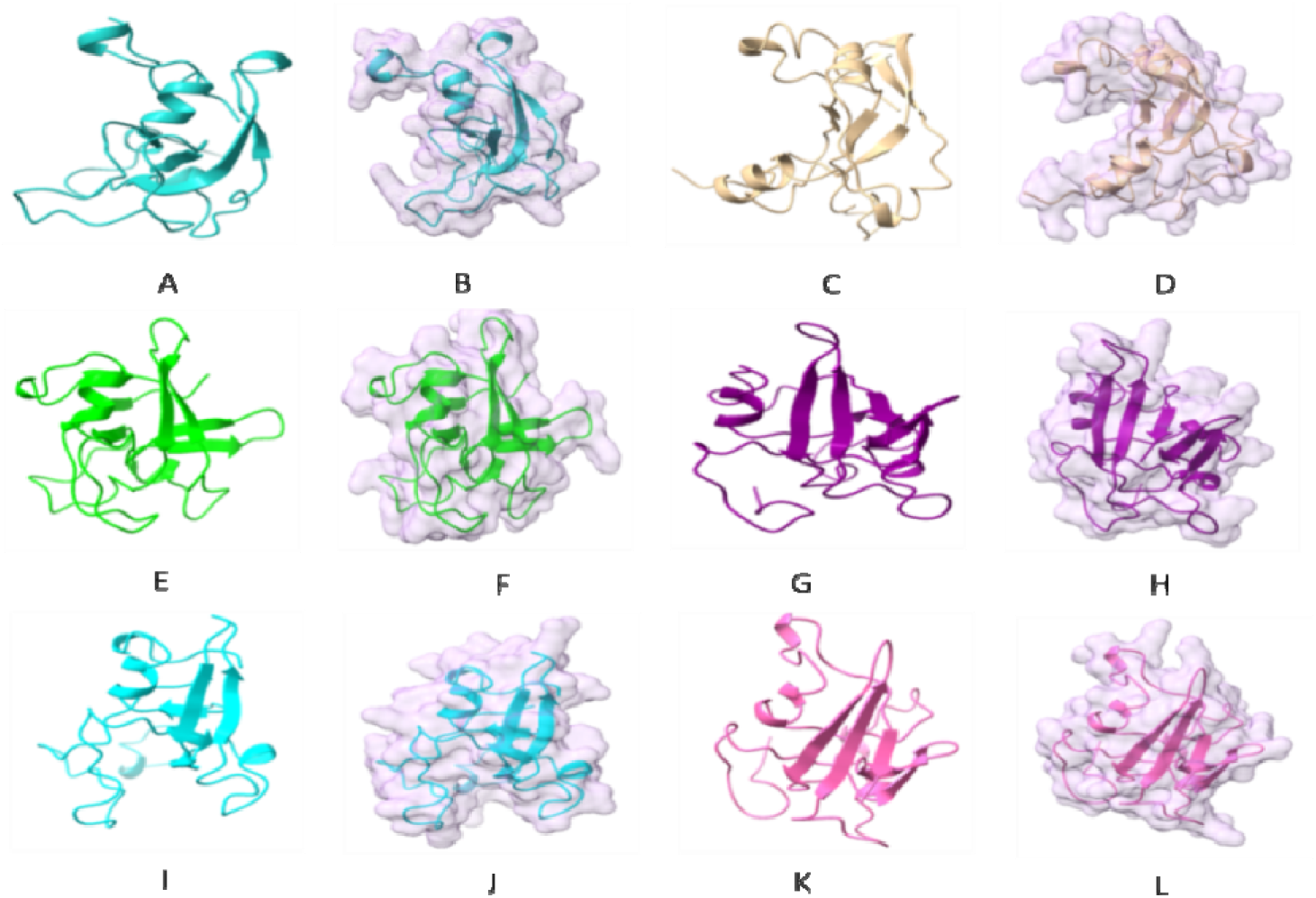
Structural modeling of selected rust kiwellin like fold containing proteins in ribbon and surface from. (A) and (B) Pstr_13960.(C) and (D) Pgr_11317. (E) and (F) Mlcri_105821. (G) and (H) Psorghi_KNZ52496.1. (I) and (J) Pcoro_02952. (K) and (L) Ptri_09007.

**Fig. 4.**
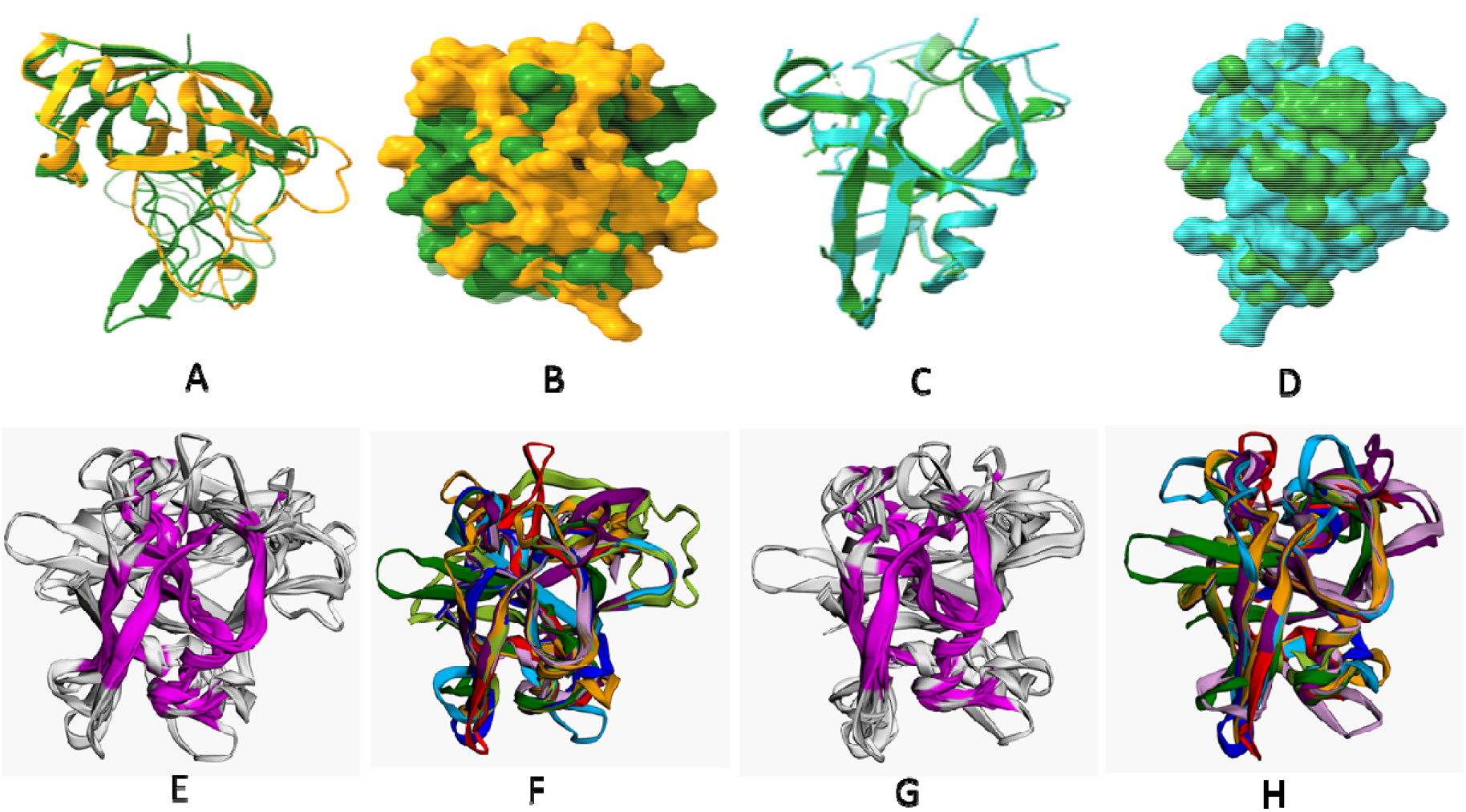
Superimposition of the selected rust KLEs and plant kiwellin proteins. (A) and (B) *Pcoro_8407* (yellow) and maize Kiwellin protein, 6TI2 (green) superimposition in ribbon and surface from. (RMSD score-1.76, Tm score-0.6). (C) and (D) Psorghi_KNZ52496.1(Cyan) and maize Kiwellin protein, 6TI2 (green) superimposition in ribbon and surface from, (RMSD score-1.76, Tm score-0.6). (E) and (F). The violet region is the core conserved region for all superimposed proteins and superimposition of Psorghi_KNZ52496.(blue), Mlcri_105821(green), Malli_1608120(red), Pcoro_02952 (magenta), Pgr_11317 (orange), Pstr_13960(sky blue), Pustr_104E_500 (violet) and 6TI2 (olive green), (ccRMSD-1.02). (G) and (H) The violet region is the core conserved region for all superimposed protein and superimposition of Psorghi_KNZ52496.(blue), Mlcri_105821(green), Malli_1608120(red), Pcoro_02952(magenta), Pstr_13960(sky blue), Pustr_104E_500( violet) and Ptri_09007 (orange) (ccRMSD −1.06)

### Intron-exon organization analysis showed rust kiwellin effector members to be evolutionary conserved despite sequence diversity

Multiple sequence alignment of the fungal kiwellin fold-like proteins showed diversity in their sequence (Supplementary Fig S1). Despite the diversity, these proteins tend to have a core kiwellin matching region that was significantly conserved when compared to full proteins (Supplementary Fig. S1. The analysis of intron-exon boundaries of randomly selected KLEsfrom *P. striiformis* showed that they have a similar intron-exon organization (Supplementary Fig S2). These randomly selected rust genes grouped into two categories where one group had nine exons and the other had eight exons. Though the size of the exons and intron more or less remained the same. Further, the alignment of full fungal kiwellin proteins with plant kiwellins also showed these proteins to be quite diverse from the later one, except at the kiwellin domain region (Fig. 1). Intriguingly, the fungal kiwellin like proteins matched at the c-terminal of plant kiwellin sequences rather than the full region.

### Similarities and differences among rust KLEs and plant kiwellins members

The phylogenetic analysis of rust KLEs using Maximum likelihood (ML) methods showed that these proteins could be classified into ten major subgroups. (Fig. 5) Further, comparative phylogeny of rust KLE proteins with plant kiwellin proteins showed grouping into five major groups distinguishing plant and fungal members (Fig. 6). However, few rust KLE proteins were found to be clustered with plants kiwellin proteins, signifying more similarity of these rust KLE members to the plant, and further suggesting an evolutionary relationship of these proteins (Fig. 6).

**Fig. 5.**
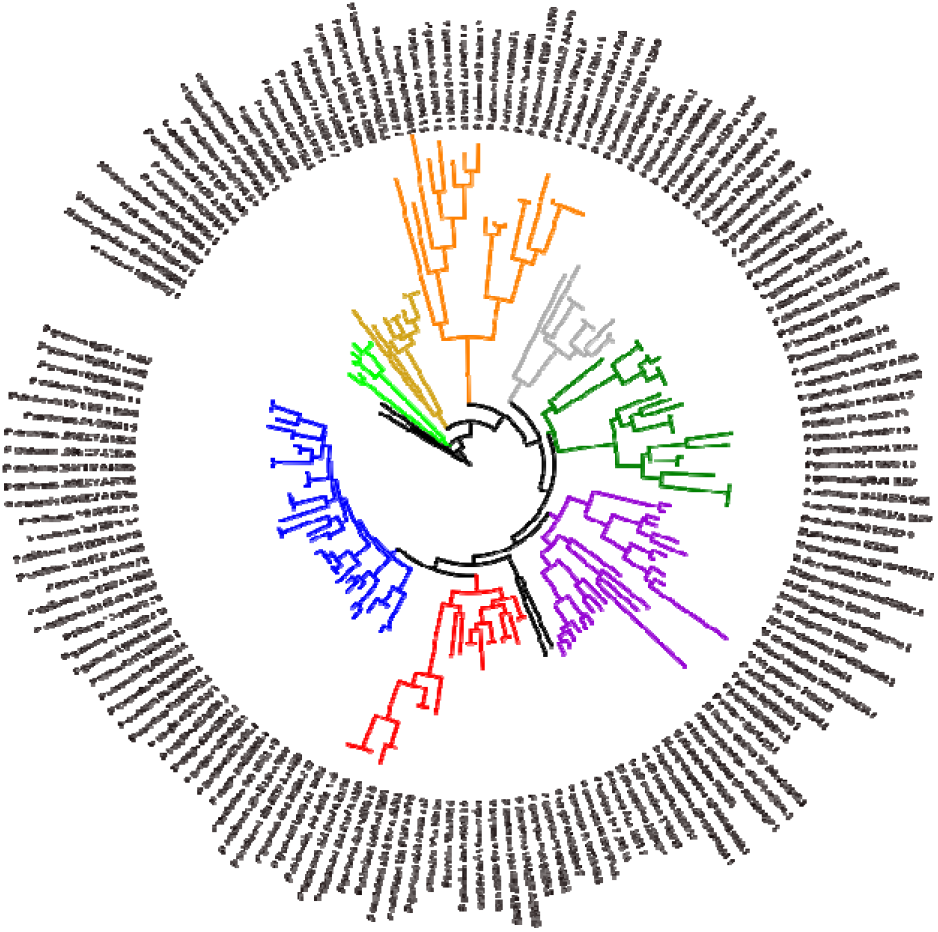
Phylogenetic analysis of rust KLEs proteins. A. Phylogenetic analysis of rust KLEs classified these proteins into ten different groups.

**Fig.6.**
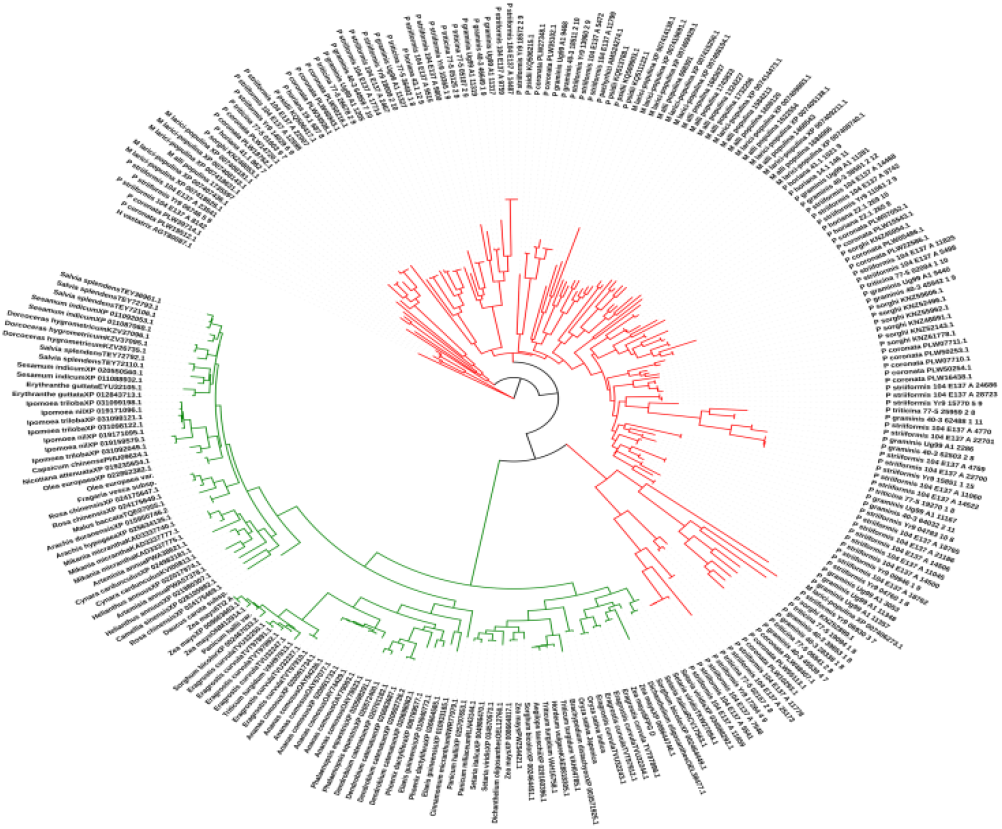
Phylogenetic analysis of rust KLE proteins and plant Kiwellin proteins. Selected plant kiwellin proteins (model plants species) were used for comparing the kiwellin proteins. The rust and plant proteins formed separate clusters but few rust KLEs clustered more closely to plant kiwellins. Plants kiwellins (green), rust KLEs (red).

### Kiwellin protein-coding genes are present in the majority of eukaryotic plant pathogens

The identification of KLEs in other plant pathogens using rust proteins as a query did not give any significant results however using plant kiwellins proteins as a query the search gave 98 hits belonging to different fungal species (Supplementary file S15, Maximum hits selected 500). We used these proteins for structural modeling to find any significant similarity at the 3D level to kiwellin proteins. Out of 97, 39 proteins showed Kiwelin like-fold at the 3D structure level (Supplementary file S16). Further, the secretory and effector analysis returned only one effector candidate (*Tremella mesentrica*, RXK38439.1) (Supplementary file S17).

### Most of the KLEs including *Pstr_13960* were predicted with N-terminal intrinsically disordered regions (IDRs) of variable length

During sequence alignment analysis, it was interesting to note that most of the fungal kiwellin fold-like proteins matched with plant kiwellin proteins at the C-terminal than the full region. The further analysis of the unmatched region Rust KLEs showed most of these proteins to possess a significantly long Intrinsically disordered region at the N-terminal (>20 % of the full length of protein) Fig.7, (Supplementary file S18).

**Fig. 7.**
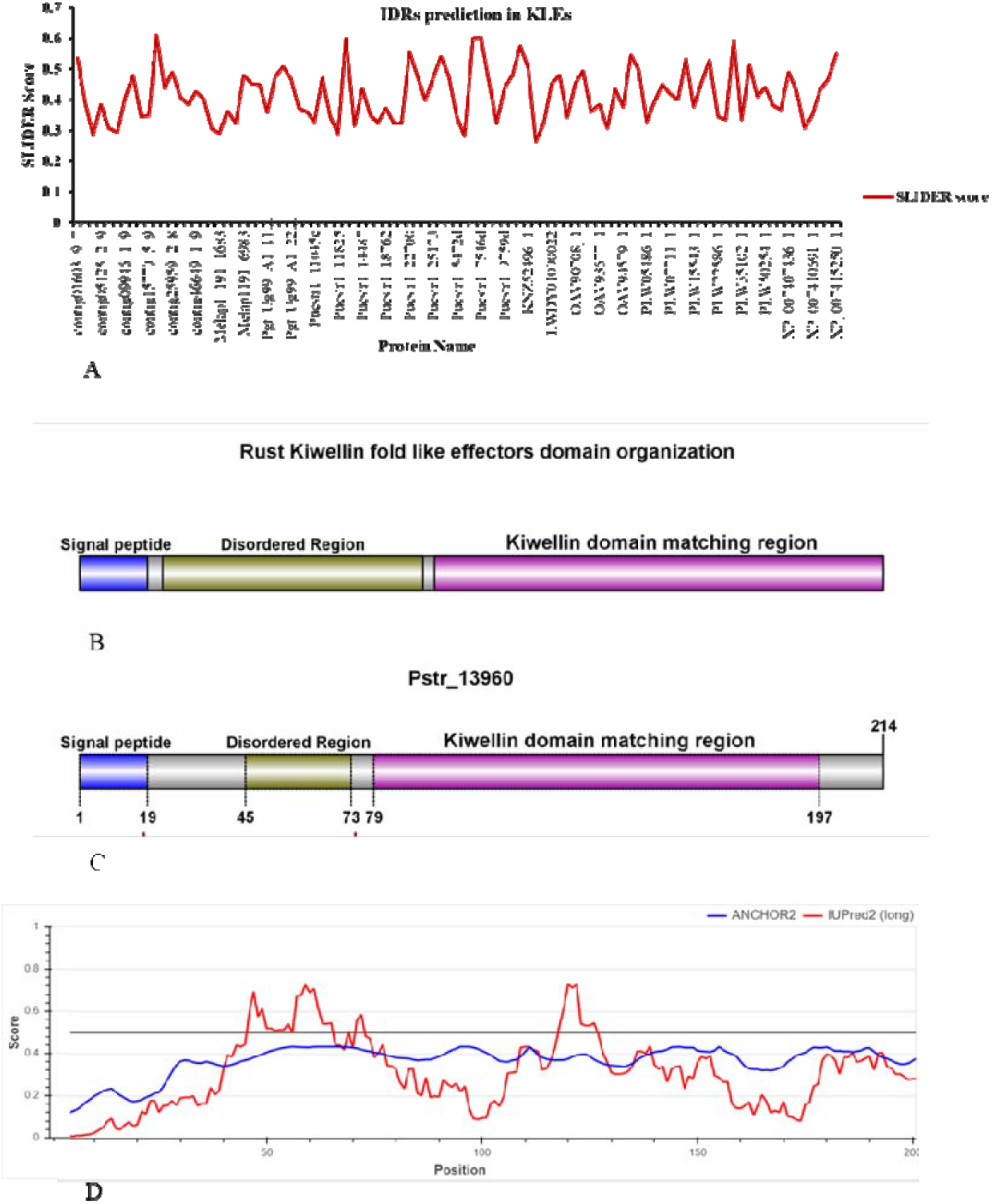
Disorder prediction and domain organization of Rust KLEs. (A) Disordered region prediction in selected rust KLEs using SLIDER software. (B) General rust KLEs domain organization (c) *Pstr_13960* domain organization (D) Intrinsic disorder region prediction of *Pstr_13960* using IUPred2A software.

The IDRs analysis using SLIDR software predicted these proteins to possess IDRs by returning different scores for proteins. The longer the IDRs the higher the score. Most of the rust KLEs were in the size of (185 to 360 amino acids) therefore the length of IDRs was also manually analyzed for the significant disordered region prediction using PONDR disorder prediction and IUPred2A.

### *Pstr_13960*, a candidate rust Kiwellin effector suppresses BAX induced cell death and localizes to the chloroplast in *N. benthamiana*

The Pstr_13960 showed the highest modeling confidence and 30% sequence similarity therefore was selected for functional identification. The transient overexpression studies of *Pst_13960* in *N. benthamiana* using agroinfiltration did not show any cell death-inducing phenotype; however, the *Pst_13960* was able to suppress BAX-induced cell death in *N. benthamiana* leaves when BAX was agroinfiltrated 24hrs after *Pstr_13960*. Further, the subcellular localization analysis in *N. benthamiana* showed *Pstr_13960* to be primarily found to localize in the chloroplast (Fig.8)

**Fig 8.**
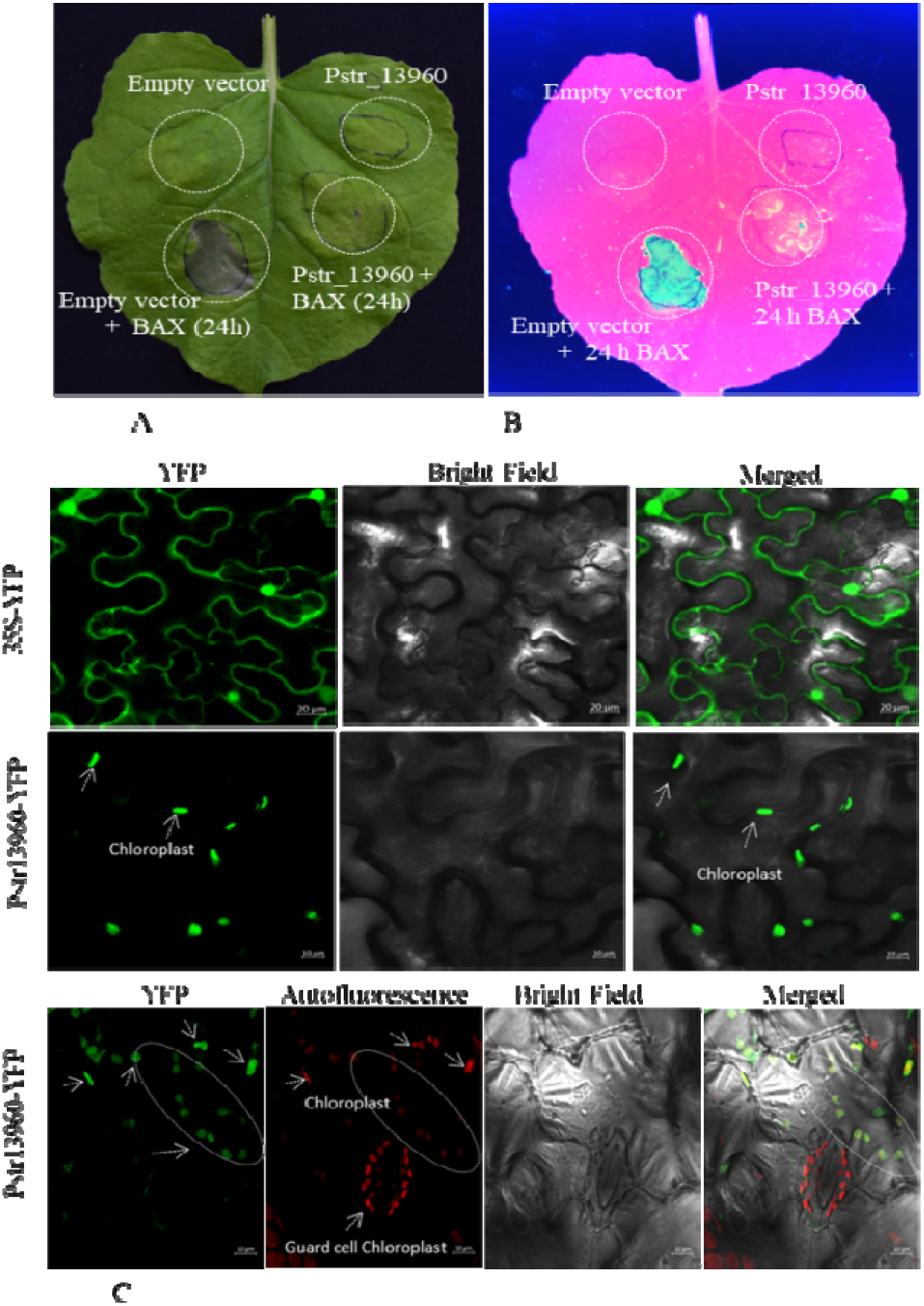
BAX induced cell death suppression assay and localization of *PStr_13960*. (A) N.benthaminan leaf showing *Pstr_13960* suppresses the BAX-induced cell death. (B) The leaf as in figure A under image analyzer (RGB color) (c) Localization of *Pstr_13960* in *N. benthamiana* using agroinfiltration showed its location in chloroplasts.

### N-terminal region near Intrinsically disorder region putatively contain chloroplast targeting region of *Pstr_13960*

To identify the chloroplast targeting region of *PStr_13960* various deletion fragments were created (Fig.9A). The full Pstr_13960 gene encoding protein primarily localizes to chloroplast. The removal of the signal peptide encoding region also did not affect the chloroplastic location of *Pstr_13960*. Further, the deletion of the Kiwellin encoding region also did no change the location of *Pstr_13960*. Surprisingly the expression of the kiwellin matching region alone after the deletion of the N-terminal region and IDR showed it to primarily localized to cytoplasm and nucleus. (Fig.9B).

**Fig 9.**
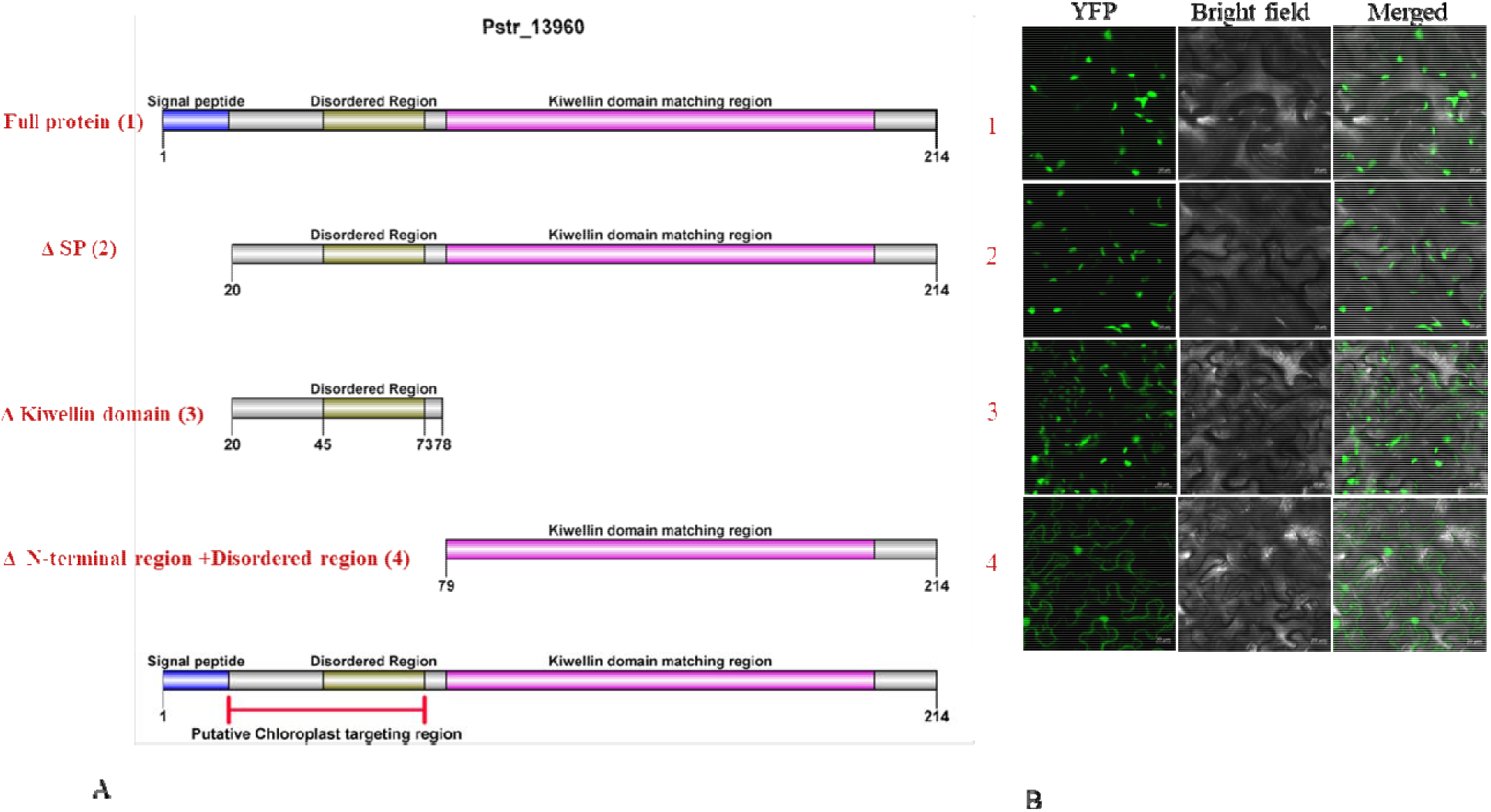
Various deletion fragments of *Pstr_13960* regions targeting chloroplast, plasma membrane, and nucleus in *N. benthaminana*. (A) Representation of deletion of various regions of *Pstr_13960* for analyzing the chloroplast targeting region. (B) Subcellular localization analysis after various deletions. A region near signal peptide and disordered region putatively encode chloroplast targeting region as kiwellin matching region alone localized to cytoplasm and nucleus. 1. Full protein, chloroplast. 2. Without signal peptide, chloroplast. 3. Without kiwellin region, chloroplast. 4. Kiwellin region only, cytoplasm and nucleus.

### The kiwellin matching region of *Pstr_13960* alone is sufficient to suppress the BAX induced cell death

To further characterize the kiwellin matching region *Pst_13960_Kiwi* was also expressed in *N. benthamiana* using BAX cell death suppression assay. The *Pst_13960_Kiwi* also suppressed the BAX-induced cell death however its location was changed from chloroplast to the nucleus and cytoplasm (Fig 10).

**Fig 10.**
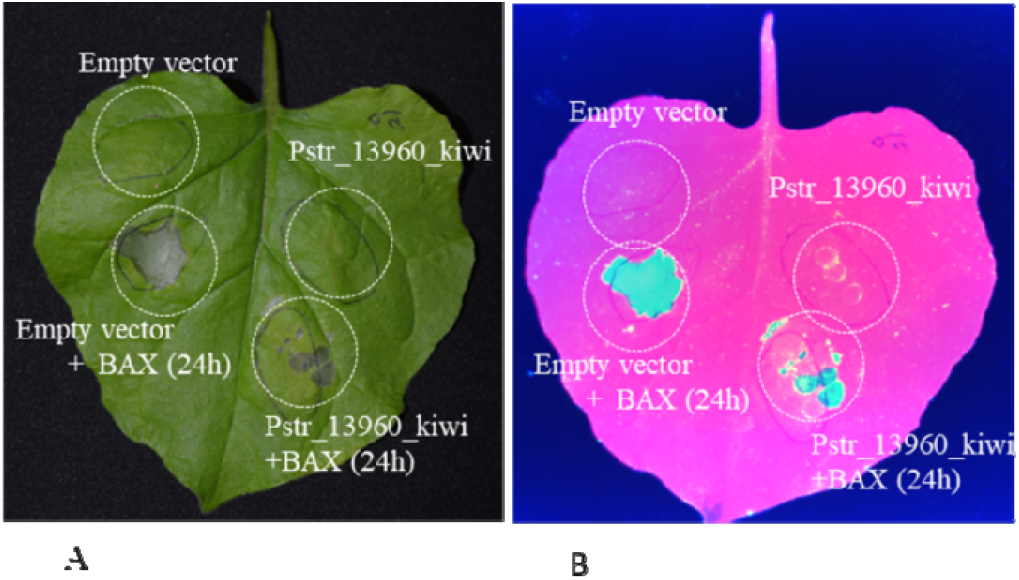
Cell death suppression assay for *PStr_13960*_kiwi using agroinfiltration. **(A)** *N. benthamiana* leaf showing BAX induced cell death suppression. (B) The leaf is in figure A under the image analyzer (RGB color).

### Interaction of rust kiwellin like effectors with plant chorismate mutase proteins (CMs) proteins

The Protein-protein docking of the maize, wheat, tobacco with rust Kiwellin proteins showed these effectors may bind with the plant CM proteins rust kiwellin like effector *Pstr_13960*. The interaction performed with all the 5 different rust effectors protein with different CMs proteins of maize, tobacco, wheat, showed positive binding (Fig. 11).

**Fig 11.**
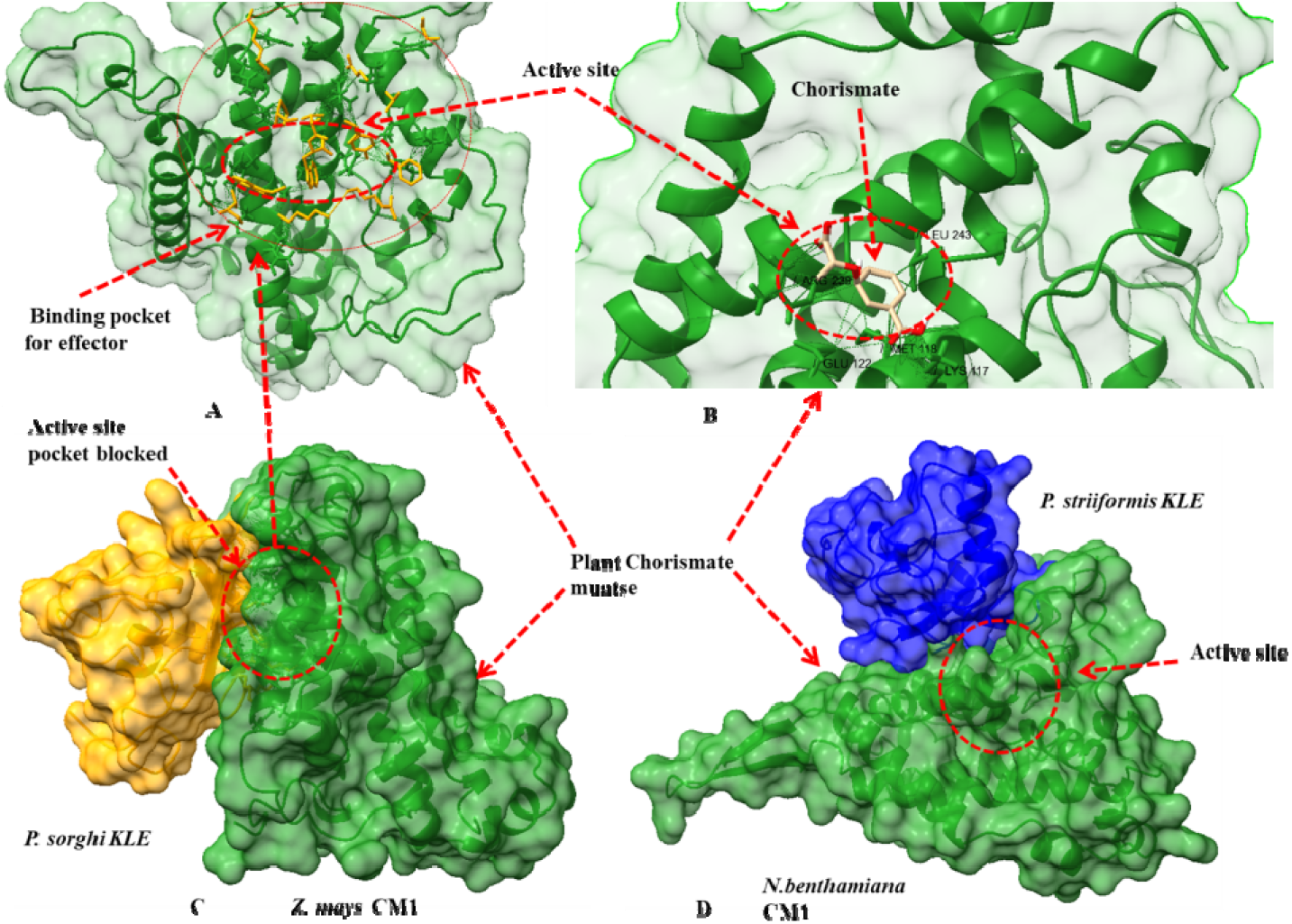
Molecular docking analysis of rust KLEs with plant Chorismate mutases. (A)Interaction of Psorghi_KNZ52496 effector with maize chorismate protein (CM1) showing binding pocket. (B) Binding of chorismate in the maize CM1 pocket. (C) binding of Psorghi_KNZ52496 covers the chorismate binding pocket of maize CM1. (D) binding of Pstr_13960 covers the chorismate binding pocket of *N. benthamiana* CM1.

## Discussion

Rust fungi are currently the most complex obligate biotrophic pathogen among all the plant pathogens. They are known to encode a large number of secretory proteins that play a major role in the host evasion and infection process explicitly known as effector proteins. Characterization of the rust effector proteins has always remained a challenge due to their species-specific nature, unique strategies by the pathogen to disarm the host, and no stable genetic transformation methods for these pathogens. Recently novel strategies implemented by effectors especially rust fungi effectors have been highlighted in various studies such as suppressing host RNA interference (RNAi), targeting various cellular organelles to suppress host defense systems, and utilizing small conserved folds to performing various biological processes. The identification of the small conserved folds in the rust effectors proves to be more useful as rust effector protein-encoding genes tend to change their sequence frequently due to the selection pressure imposed by host Resistance genes (R). Considering this fact, in the present study we have performed comparative secretome and computational structural genomic studies to identify novel conserved folds in plant-pathogen including rust fungi. The identified unannotated secretory proteins showed several variations in the studied species. The biotrophs followed by hemibiotrophs generally encode a large number of novel proteins. However, the necrotrophs require the least amount of novel small secretory/effector proteins as these pathogens use mostly cell wall degrading enzymes and toxins to complete the infection cycle was also reflected in our study. Further, the novel unannotated secretome of plant pathogens rarely share any similarities at the primary sequence level, as well as at a structural level, was also consistent in our results, as most of these proteins did not show any similarity to conserved fold at the threshold (>20% sequence similarity, 60% modeling confidence). However, in the rust secretome, we found few proteins (1-4) per species that were matching to the maize and kiwi kiwellin proteins. The kiwellin proteins are reported to be present in all major plant species including kiwi fruit, where these proteins are regarded as a major allergen. Though, recent studies have reported the functioning of these proteins in plant-pathogen interaction [20]. In potatoes, the protein-coding genes of these proteins were highly upregulated when infected with *P. infestans*, suggesting the crucial role of these proteins in host defense[39,40]. Moreover, a major role of these proteins was highlighted as these were predicted to bind to major effector chorismate mutase (*Cmu1*) in the fungal pathogen that is known to affect salicylic acid production in chloroplast to alter the defense system of the plant. In this study, Han et al., 2019 showed that the plant encodes a large number of kiwellin proteins where many of them could be specific to pathogen CMs rather than the host-encoded CMs[20]. The presence of kiwellin like secretory proteins in rust pathogens was very surprising and interesting to see as none of the studies has reported these folds to be present in any other species. Although the study carried out by Han et al.,2019 also reported the presence of kiwellin like proteins in eukaryotes including various fungal species, no detailed study was carried out to analyze their function as an effector or virulence factor[20]. In our study, we found KLEs in rust alone. Therefore, to rule out the possibility, the identification of the kiwellin protein fold in other plant pathogens showed these proteins are also present in diverse plant pathogens with significant conservation. Out of 97 proteins matching with maize kiwellin sequences, the 39 proteins showed conserved fold similarity to maize kiwellin, but only one protein showed (*Tremella mesentrica*, RXK38439.1) the signature of signal peptide and effector-like feature (Supplementary file S17 and S18). In contrast, the majority of the identified candidates in rust (Table2) were predicted secretory and effector candidates when analyzed. These results suggested that rusts have evolved these proteins as a potential effector family as a high number of KLEs was present in most of the studied rust species (maximum 28 proteins). The kiwellin proteins are known to possess a DPBB domain that matches with several other protein families with expansion-like features, sugar-binding proteins[20,21,23]. Though structural modeling significantly distinguished proteins with kiwellin like fold and sugar-binding fold-containing proteins. The results further highlighted the fact the structural modeling can distinguish between homologous proteins however with the different folds at the 3-dimensional level.

**Table 2.** Secretory and effector feature analysis of Rust kiwellin like effector of rust fungi.

| Organism Name | Proteins matching with<br>Kiwellin proteins like<br>fold* | Secretory<br>Proteins | Proteins with effector<br>like features<br>(Effector P 2.0) |
| --- | --- | --- | --- |
| *Puccinia graminis* UG99 A | 12 | 8 | 8 |
| *Puccinia graminis* 40-3 | 12 | 5 | 3 |
| *Puccinia triticina* 77-5 | 11 | 6 | 5 |
| *Puccinia triticina* Race1 | 16 | 11 | 10 |
| *Puccinia striiformis* Yr9 | 14 | 9 | 8 |
| *Puccinia striiformis* 104 E137 A | 35 | 28 | 26 |
| *Puccinia coronata* | 21 | 18 | 17 |
| *Melampsora larici-populina* | 15 | 11 | 8 |
| *Melampsora allii-populina* | 10 | 7 | 6 |
| *Puccinia horiana* | 7 | 3 | 2 |
| *Puccinia sorghi* | 9 | 3 | 3 |
| *Puccinia psidii* | 3 | 1 | 1 |
| *Phakopsora pachyrhizi* | 1 | 1 | 1 |
| *Hemileia vastatrix* | 1 | 1 | 0 |

\*Novel as well as with DPBB domain matching proteins

The presence of a variable number of KLEs in rust fungi could be explained by two possibilities. First, these species have been infecting the host in different geographical locations, as a result, may have expanded the gene family due to variable selection pressure by the host species. The second possibility is the use of different sequencing platforms for sequencing the rust genomes as a result of a partially assembled genome. The highest number of KLEs were found in two recently sequenced genomes *P. striiformis 104E137A* and *P. coronata*, where both of them have been sequenced by long-read sequencing technologies [41,42]. The fewer KLEs found in all other previously sequenced genome assemblies could be the result of short-read sequencing technologies or incomplete assemblies of these genomes. Considering the results of two advanced read technologies assemblies we could speculate that the rust genome may encode several KLEs, however, many of them could not be identified due to short read fragmented assemblies of the other rust genomes.

The structural analysis of the rust KLEs showed these proteins to possess a similar fold as that of plant kiwelin protein predicting the rust KLEs to be involved in a similar biological process (Fig. 3). However, the structure was not fully identical as the rust proteins may have a diverged function than original plant kiwellin proteins as suggested by superimposition studies. Further, the intron-exon organization studies of randomly selected rust candidates showed overall similar organizations suggesting these members have a common origin. Though one of the groups possesses one additional exon proposing the further evolution of the members to attain novel function among themselves. The phylogenetic analysis of the rust KLEs classified these proteins into ten groups describing the diversity of these proteins. Moreover, the diversity in these sequences could also be supported by the structural modeling results, as these proteins showed variable modeling confidence and sequence similarity to plant kiwellins. The two distinct clades formed in plant and rust kiwellins showed these proteins to be independently involved, though the similarity of few rust members with plant kiwellins further suggested some of these proteins may perform relatively similar functions to plant kiwellins.

The presence of the IDRs in the proteins is predicted to impart these proteins diverse functions. In the case of the bacterial effector proteins, it is associated with several functions such as translocation, secretion, interaction with multiple candidates. In the case of oomycetes, the presence of IDR is a common structural characteristic for several RxLR and WY effectors proteins[43,44]. The RxLR region is also known to act as a universal translocation motif in a large number of oomycetes proteins. The study carried out by Shen et al.,2017 also showed RxLR predicted region in these oomycetes effectors is preferably disordered[45]. Moreover, the mutation in the disordered region also leads to loss of avirulence activity for PsAvh18 and PcAvh207 effectors further suggesting the importance of the disordered region in effectors proteins[45]. Recently, a study on RxLR effector Avr2 showed that IDR is important for its effector function [46]. The presence of IDRs in the rust KLEs at N-terminal also suggested them to encode putative transient peptides or some other crucial function as the subcellular localization studies showed one of the *P. striiformis* KLE *Pstr_13960* to located in the chloroplast. The N-terminal region or the signal peptide region alone could be associated with the translocation was ruled out as removal of signal peptide encoding region of the *Pstr_13960* also localized it on the chloroplast. Further N-terminal region (after signal peptide and before predicted IDR) was also very small (25 amino acids) further supporting the role of IDR in translocation as the average length for plastid transient peptide region is 50 amino acids in most of the studies[47] (Fig. 9). The role of IDR and N-terminal region in chloroplast translocation was revealed as the deletion of this region localizes the *Pstr_13960* near the membrane and nucleus.

The BAX-induced cell death suppression activity of *PStr_13960* showed its potential role as an effector. Surprisingly, the expression of the Kiwellin predicted region alone further suggested that the kiwellin matching region is responsible for cell death suppression activity of *Pstr_13960* signifying the novel role of Kiwellin like fold proteins in rust fungi. As kiwellin proteins are reported to play important role in plant defense, the utilization of kiwellin like fold to disarm the host itself uncovers the novel strategies of pathogens. The kiwelin proteins in plants are known to stop the activity of the *Cmu1* effector by inhibiting its binding to host chorismate proteins so that enough salicylic acid (SA) could be produced to defend against pathogen infection especially biotrophs. As the *Cmu1* is reported to present in diverse fungal pathogens this mechanism is applicable in most of the plant pathogens interactions. We also identified CM effectors in rust fungal pathogens using the NCBI NR database as well as using BLASTP search against individual rust proteins set. We found one to two copies of secretory Cmu in *Puccinia* species in the rust genome. The SA plays an indispensable role in the host defense system against all biotrophic pathogens including rust fungi. In this scenario, the role of rust KLEs could be highlighted. The function of the rust effector could be highlight by the alternate strategy of rust fungi to halt SA synthesis. As plant kiwellin binds to fungal CMs, the rust KLEs may also follow the same strategies of binding plant CMs proteins to affect the SA synthesis.

As plant kiwellin binds to fungal CMs allowing plant CMs to produce SA synthesis, in response to this, as a counter-strategy the kiwellin effectors may bind to free plant CMs to suppress SA production and promote infection. As an alternate strategy, the rust fungi may have evolved this strategy in the course of evolution to fail kiwellin mediated defense response. Moreover, as plant chorismate mutase primarily localizes in chloroplast and few of them localizes to the cytoplasm, the *Pstr_13960* and *Pstr_13960_kiwi* also show the same location as predicted in subcellular localization studies suggesting the interaction of *Pstr_13960* with plant CMs is feasible. Additionally, the docking analysis also showed rust KLEs can interact to plant CMs in docking studies further strengthen the possibility of interaction *Pstr_13960* with plant CMs. The residues involved in the CM interaction are also conserved in the rust KLEs when compared with maize KWL1 and KWL1b (Fig. 2). Though, further interaction or colocalization studies may validate the interaction of rust KLEs with plant CMs.

## Conclusions

The kiwellin proteins have been known recently for the plant defense function against pathogens, however, these proteins are present in several eukaryotic species. In this study, we report the presence of these proteins in the plant-pathogen genome and showed them to act as effectors in rust fungi. We propose that kiwellin like proteins are widely present in eukaryotes including plant pathogens, however, selected species such as rusts have evolved, expanded them, and to use as effector proteins by translocating to the chloroplast and suppressing plant cell death. We also predicted that rust KLEs are acting as effectors probably by interacting to plant CMs as a counter-strategy when host kiwellin proteins bind to rust CMs. The evolution of kiwellin proteins as effectors could be alternate strategies of rust fungi to avoid plant kiwellin proteins mediated host defense when chorismate mutase effector mediated host manipulation mechanism becomes ineffective.

## Data Availability Statement

The datasets analyzed in this study can be found publically (https://github.com/rajdeepjaswal52/Kiwellin-proteins.git).

## Supporting Information

**S1 Fig.1 Multiple sequence alignment of identified rust KLE proteins.** 109 identified rust KLEs were variable in length up to 360 amino acids, therefore to completely highlight the aligned region the N-terminal region was showed in this figure (PDF).

**S2 Fig. Intron-exon analysis of selected rust KLE encoding genes.** Randomly, 12 genes were selected from *P. striiformis race 104E* and analyzed for the conserved pattern. The size of the exon more or less remained the same for these genes however few of the genes had one additional exon suggesting the evolution of the family (PDF).

## Supplementary Excel file

**S1.** Structural modeling of the novel kiwellin matching effectors Identified in this study initially

**S2**. DPBB domain matching proteins in the rust secretome

**S3**. Structural modeling of DPBB domain matching proteins in the rust secretome.

**S4-S15**. Effector prediction of the kiwelin fold containing proteins in the secretome of the different species rust.

**S16.** Structural modeling of DPBB/kiwellin domain matching proteins present in plant pathogens other than rust fungi.

**S17**. Secretory and effector prediction of DPBB/kiwellin domain matching proteins in plant pathogens other than rust fungi.

**S18**. Intrinsic disorder prediction in the rust KLE protein sequences.

## Acknowledgments

TRS is thankful to the Department of Science and Technology, Govt. of India, for JC Bose National Fellowship. RJ is thankful to the University Grants Commission (UGC), New Delhi for providing Junior Research Fellowship (JRF). The authors also gratefully acknowledge Miss Aakriti Mehra for assistance in confocal imaging experiments.

## Author Contributions

**Conceptualization:** Tilak Raj Sharma and Rajdeep Jaswal

**Data curation and lab experiments:** Rajdeep Jaswal

**Data analysis and writng first draft:** Rajdeep Jaswal, Sivasubramanian Rajarammohan, Himanshu Dubey, Kanti Kiran, Hukam Rawal, Humira Sonah, Rupesh Deshmukh

**Funding acquisition:** Tilak Raj Sharma

**Investigation:** Rajdeep Jaswal

**Methodology and writing final draft:** Tilak Raj Sharma and Rajdeep Jaswal

