## Supplementary figures and images for "A kiwellin protein-like fold containing rust effector protein localizes to chloroplast and suppress cell death in plants"

### Fig. S1_ rust_kiwellin_alignmnet_full_compressed.pdf

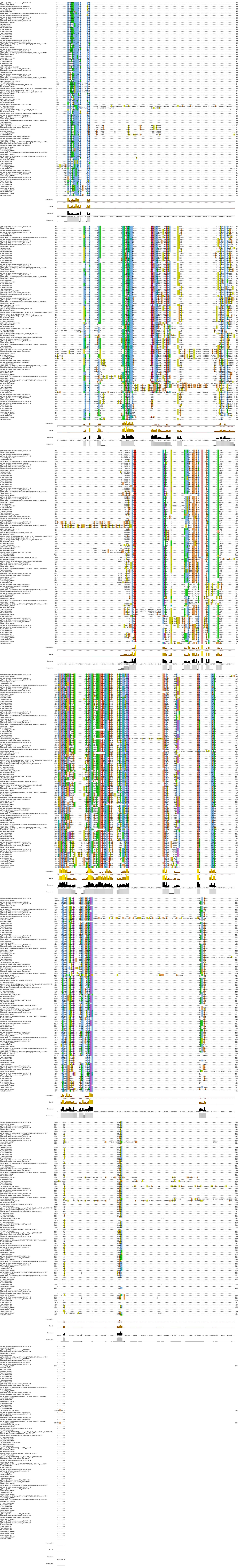

### FIg. S2_Intron_exon analysis_compressed.pdf

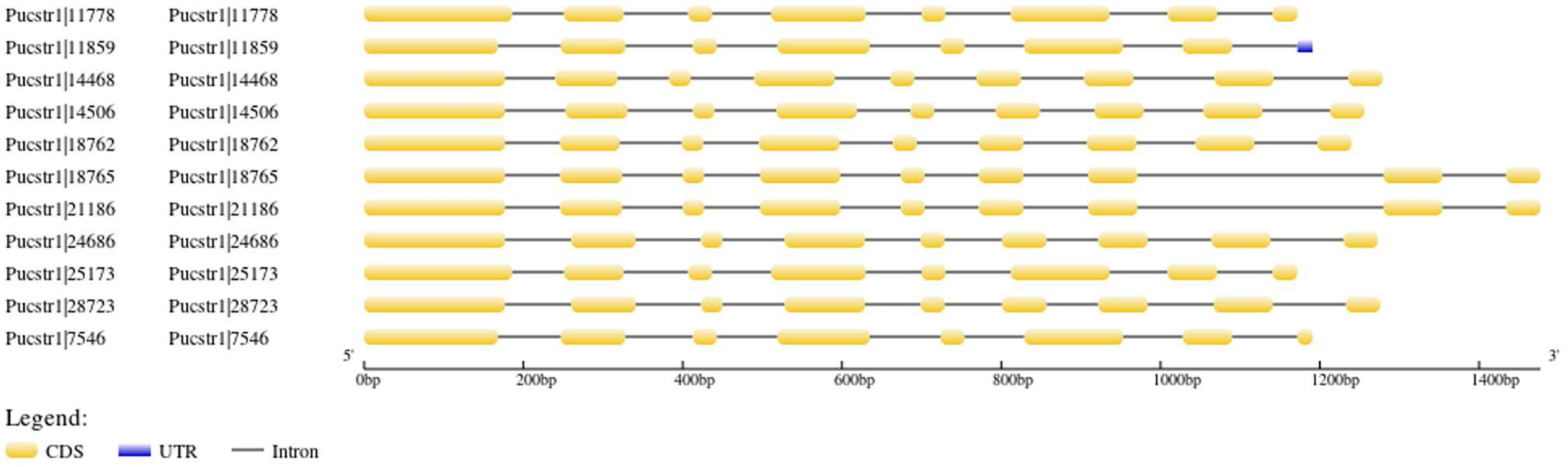
